# Integrative Comparison of GeneHancer and Single-Cell Co-accessibility Reveals Active Enhancer–Gene Interactions

**DOI:** 10.64898/2026.01.23.701222

**Authors:** Lorenzo Martini, Roberta Bardini, Alessandro Savino, Stefano Di Carlo

**Affiliations:** Politecnico di Torino

**Keywords:** Multi-omic, Single-cell, scATAC-seq, Transcriptional Regulation, GeneHancer, Co-accessibility

## Abstract

Linking enhancers to their target genes remains challenging due to the context-independent nature of curated annotations and the noise inherent in data-driven predictions. GeneHancer provides a comprehensive catalogue of enhancer–gene associations, but many elements are inactive in specific biological settings. Conversely, co-accessibility inferred from single-cell chromatin accessibility data captures sample-specific regulatory structure but may reflect indirect or non-functional interactions. This work integrates these complementary perspectives by comparing GeneHancer annotations with co-accessibility networks derived from a human PBMC Multiome dataset. Using Circe to infer peak–peak co-accessibility and GRAIGH to map peaks onto GeneHancer elements, this approach identifies enhancer–gene associations supported both by prior evidence and by accessibility patterns in the dataset. Only a small subset of GeneHancer links is validated by co-accessibility, yet these conserved associations display substantially higher cell-type specificity and stronger accessibility–expression concordance than either the full or “Elite” GeneHancer sets. This refined subset isolates regulatory interactions that are both biologically plausible and active in the sample, reducing redundancy and improving interpretability. Our results show that integrating curated enhancer annotations with single-cell epigenomic evidence yields a focused, high-confidence regulatory map suited for analyzing transcriptional regulation and cell identity in a dataset-specific manner.

## 1 INTRODUCTION

The genome encodes the information underlying nearly all cellular functions, yet our current understanding captures only a fraction of its regulatory complexity [Altschuler and Wu, 2010]. While many genes are now well annotated, large portions of the non-coding genome, particularly regulatory regions, remain poorly characterized. These regions, although not necessarily transcribed into specific RNAs, are fundamental for proper transcriptional control and higher-order regulatory mechanisms [Moosavi et al., 2016]. Among these regulatory elements, enhancers constitute a particularly important class of cis-regulatory DNA sequences. Distributed throughout the genome, enhancers modulate the transcription of proximal or distally located genes by recruiting specific Transcription Factor (TF)s [Martini et al., 2024]. A substantial body of literature highlights their essential roles in cellular differentiation, development, and disease progression, including cancer [Shailendra S, 2021, Buen-rostro et al., 2018].

Despite their biological importance, enhancer definition and identification remain challenging. In contrast to protein-coding regions, enhancer boundaries and activities are often context-dependent and poorly delineated [Pennacchio and Bothers, 2013]. Establishing reliable links between enhancers and their target genes is even more difficult and typically requires extensive experimental validation or integrative computational evidence [He et al., 2014]. Existing approaches span curated databases and data-driven analyses, yet each presents substantial limitations: curated resources may include redundant, overly broad, or non-specific associations, while computational predictions are highly dependent on the underlying datasets and vulnerable to noise. In parallel, a growing number of methods aim to infer cis-regulatory and gene regulatory networks to model gene activity [Martini et al., 2023a, Kamimoto et al., 2023]. These approaches move beyond simple correlation or co-expression of TFs by explicitly incorporating cis-regulatory genomic regions. Their success underscores the need for integrative strategies that effectively combine curated biological knowledge with data-driven, context-specific evidence.

To address these challenges, this study integrates enhancer–gene associations from the GeneHancer database [Fishilevich et al., 2017] with genomic connections inferred through co-accessibility analyses applied to specific epigenomic datasets. Rather than performing a direct comparison between curated and inferred interactions, our approach leverages co-accessibility information to refine the broad and heterogeneous GeneHancer catalogue, producing a more precise and sample-relevant set of regulatory relationships. By intersecting cross-validated enhancer annotations with dataset-specific chromatin accessibility patterns, we systematically reduce redundancy in GeneHancer and prioritize enhancer–gene links that are supported by the data.

Our results demonstrate that only 2.4% of GeneHancer associations are supported by the coaccessibility analysis, highlighting the importance of tailoring curated database information to the biological context under investigation. Notably, the refined set of interactions shows substantially increased specificity and stronger correlation with gene expression for key marker genes compared to the unfiltered database. Overall, this framework provides a targeted and reliable strategy for uncovering and analyzing dataset-specific transcriptional regulation, bridging curated knowledge and data-driven inference in enhancer–gene mapping.

## 2 BACKGROUND

A growing number of genomic resources and computational tools aim to characterize the functional landscape of regulatory elements. Large consortia such as Ensembl [Dyer et al., 2025], ENCODE [Kagda et al., 2025], and FANTOM5 [Kawaji et al., 2011] integrate diverse experimental datasets to annotate promoters, enhancers, long non-coding RNAs (lncRNAs), and microRNAs (miRNAs). While these efforts provide extensive catalogs of regulatory regions, they generally do not explicitly define functional relationships between enhancers and their target genes.

This limitation is addressed by resources such as the GeneHancer database, part of the GeneCards suite [Stelzer et al., 2016]. GeneHancer compiles genome-wide enhancer–gene and promoter–gene associations, covering approximately 18% of the human genome. Its annotations are derived from nine independent evidence sources, including expression quantitative trait loci (eQTLs), enhancer RNA (eRNA) co-expression, TF co-expression, and capture Hi-C (CHi-C). This integrative strategy yields curated, cross-validated regulatory relationships supported by functional evidence, making GeneHancer a valuable foundation for studying genome-wide regulatory architecture.

Complementary to curated regulatory annotations, epigenomic technologies such as single-cell assays for transposase-accessible chromatin sequencing (scATAC-seq) provide high-resolution measurements of chromatin accessibility at the single-cell level [Kelsey et al., 2017]. These data capture the regulatory landscape in which enhancers and promoters are active in specific cellular contexts. Because chromatin accessibility is a prerequisite for transcriptional regulation, scATAC-seq offers a powerful means to infer potential enhancer–gene relationships that are specific to a given biological state [Martini et al., 2024]. However, although scATAC-seq robustly identifies accessible regulatory elements, interpreting the functional relevance of distal accessible sites remains challenging, particularly when assigning them to their target genes [Yan et al., 2020]. This limitation directly impacts the reconstruction of enhancer–gene regulatory networks, as most enhancers act over long genomic distances. Computational strategies such as co-accessibility analysis partially address this issue by inferring putative regulatory links based on co-ordinated accessibility patterns [Pliner et al., 2018]. Nevertheless, co-accessibility predictions alone often lack specificity and therefore require integration with external regulatory annotations to improve both biological interpretability and reliability.

In this work, these limitations motivate the integration of co-accessibility–derived interactions with GeneHancer enhancer–gene annotations, with the objective of refining enhancer–gene associations to those that are both epigenomically supported and biologically plausible. Although the analysis is demonstrated on an example dataset, the proposed approach is designed as a general framework for highlighting sample-specific regulatory interactions through the integration of large, heterogeneous annotation resources with cell-type–resolved chromatin accessibility signals. As shown in the Results, intersecting GeneHancer associations with co-accessibility predictions yields a substantially smaller and more functionally coherent subset of enhancer–gene links, better reflecting the regulatory architecture active in the sample (Figure 1). Notably, the resulting associations exhibit higher specificity when considering regulatory elements linked to known marker genes and show stronger correlation with gene expression, indicating an improved ability to unravel transcriptional regulation from scATAC-seq data.

**Figure 1.**
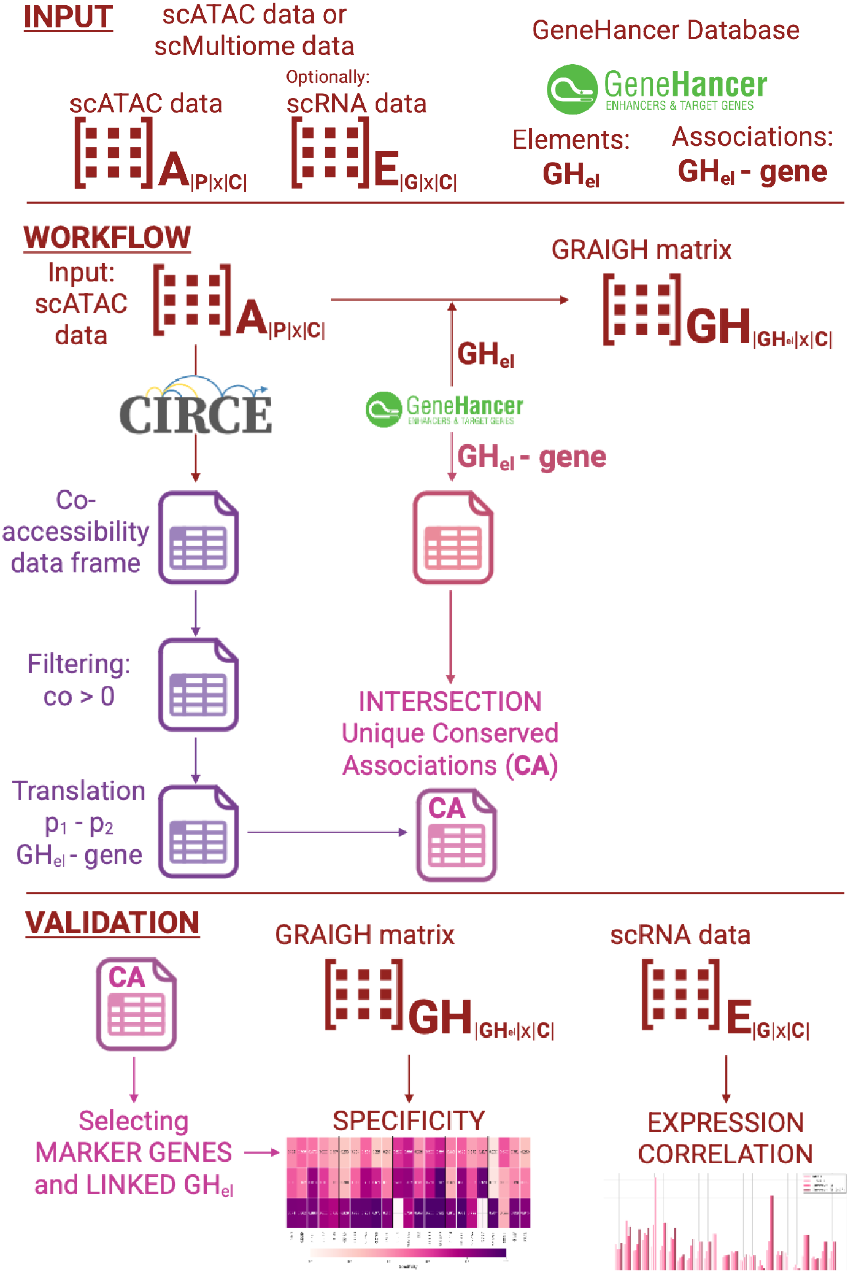
Workflow and Validation. (**1**) The input **A**_|*P*|×|*C*|_ scATAC-seq matrix is elaborated in two ways. First, it is transformed to the **GRAIGH GH**_|**GHel**|×|*C*|_ matrix. Second, the co-accessibility is calculated. Parallely, the *GH*_*el*_ ™ *g* pairs information is taken from the GeneHancer database. (**2**) The co-accessibility is further filtered and translated to be directly compared with GeneHancer. The two sets of connections are intersected to obtain the final Conserved Associations (**CA**). (**3**) The **CA** are assessed with two separated analyses employing the **GH**_|**GHel**|×|*C*|_ matrix. First, the specificity of GeneHancer element (*GH*_*el*_) linked to marker genes. Second, the correlation of *GH*_*el*_ accessibility and gene expression.

Some recent approaches also aim to characterize enhancer activity and enhancer–gene relationships using single-cell data. Among these, scEnhancer [Gao et al., 2022] provides a large-scale single-cell enhancer resource with annotations across hundreds of tissues and cell types. scEnhancer focuses on aggregating chromatin accessibility and epigenomic signals (from tens of scATAC-seq datasets) to annotate enhancer activity at scale, offering a valuable reference atlas for enhancer usage across biological contexts.

In contrast, the approach presented here is not designed to generate a global enhancer catalog, but rather to refine existing curated enhancer–gene associations within a specific dataset. By intersecting GeneHancer annotations with co-accessibility networks derived from scATAC-seq data, this framework prioritizes enhancer–gene links that are both biologically supported and active in the analyzed sample. This distinction is particularly relevant when the goal is to interpret transcriptional regulation in a defined experimental context, rather than to build a universal enhancer annotation.

Overall, the proposed framework is complementary to existing resources and methods. Large-scale enhancer atlases provide breadth across conditions, whereas this integration strategy emphasizes precision and contextual relevance, making it particularly suitable for downstream analyses of cell identity and regulatory programs in single-cell datasets.

The following section describes in detail how these associations are computed, intersected, and quantitatively evaluated against the indiscriminate use of database-derived regulatory information.

## 3 MATERIAL AND METHODS

This section describes the data sources, computational workflow, and validation strategy used to integrate curated enhancer–gene annotations with data-driven chromatin accessibility information. As illustrated in Figure 1, the proposed framework combines GeneHancer regulatory associations with co-accessibility inference from scATAC-seq data to identify a refined, dataset-specific set of cis-regulatory interactions. The workflow proceeds through three main stages: (i) transformation of chromatin accessibility into GeneHancer-centered regulatory profiles, (ii) inference and alignment of co-accessibility-derived links with curated enhancer– gene associations, and (iii) validation of the resulting conserved associations using specificity and accessibility–expression correlation analyses.

### 3.1 Input data

Two input sources are required: a scATAC-seq dataset (or, alternatively, a multiome dataset including an ATAC assay) and the GeneHancer regulatory database.

GeneHancer, part of the GeneCards suite, provides curated information on human transcriptional regulation by reporting genome-wide associations between enhancers and genes, as well as between promoters and genes. Regulatory elements are derived from a comprehensive cross-source integration of nine independent data sources, including expression quantitative trait loci (eQTLs), enhancer RNA coexpression, TF co-expression, and capture Hi-C. This integration yields reliable, non-redundant regulatory annotations supported by functional evidence. Each *GH*_*el*_ is uniquely identified by genomic coordinates, associated target genes, and a confidence score, with *Elite* associations validated across multiple independent sources. The current release includes 393,464 *GH*_*el*_ and 2,408,198 enhancer–gene connections.

As scATAC-seq data source, this paper evaluated the approach on a high-quality and well-characterized dataset, namely a 10x Genomics Multiome Human peripheral blood mononuclear cells (PBMC) dataset comprising simultaneous single-cell RNA sequencing (scRNA-seq) and scATAC-seq profiles for 10,091 cells [10XGenomics, 2018]. The dataset includes major immune cell populations, i.e., monocytes, T cells, B cells, NK cells, and rarer subtypes, which were previously annotated using RNA-based Seurat label transfer. The final annotated dataset contains 14 cell types.

This study primarily focuses on the scATAC-seq modality, represented by the accessibility matrix

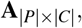

where *P* denotes peaks and *C* cells. The scRNA-seq modality, represented by

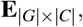

with *G* genes, is used exclusively for downstream validation analyses.

All preprocessing and analyses are implemented in Python within the *scverse* ecosystem [Virshup et al., 2023]. Multiomic data are handled using the Muon data structure [Bredikhin et al., 2022], and standard preprocessing follows the Scanpy workflow [Wolf et al., 2018].

### 3.2 Workflow

This is the core of the proposed method aiming at integrating GeneHancer regulatory elements with scATAC-seq data.

#### 3.2.1 Processing and integration of scATAC-seq data

The first step applies Gene Regulation Accessibility Integrating GeneHancer (GRAIGH) [Martini et al., 2023b] to the input accessibility matrix **A** _|*P* |*×*|*C*|_. GRAIGH maps peak-level accessibility to GeneHancer regulatory elements, generating a new matrix

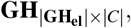

where features correspond to *GH*_*el*_ rather than dataset-specific peaks. This transformation provides two key advantages. First, *GH*_*el*_ represent uniquely defined and interoperable regulatory features, enabling direct comparison across datasets, a property that is not trivial for peak-based representations. Second, it enables direct investigation of cis-regulatory elements with established biological meaning, facilitating interpretation of cell-type–specific regulatory mechanisms. The resulting matrix is normalized using Scanpy and stored in the Muon object.

#### 3.2.2 Co-accessibility inference from scATAC-seq data

In parallel, the original accessibility matrix **A** _|*P*|*×*| *C*|_ is analyzed to infer co-accessibility between genomic regions. Co-accessibility identifies pairs of regions that tend to be accessible in the same cells, providing candidates for cis-regulatory interactions.

Co-accessibility is computed using Circe [Trimbour et al., 2025], a regression-based framework relying on graphical LASSO [Friedman et al., 2008]. To ensure computational feasibility and biological relevance, co-accessibility is calculated only between peaks within a predefined genomic distance. This distance determines both the maximum interaction range and the distance-dependent penalty used in the model. Following Circe guidelines, a maximum distance of 5 Mbp is employed, which also aligns well with the genomic span of most GeneHancer enhancer–gene associations.

Circe supports both single-cell and metacell-based computation. In this study, metacells are used to mitigate the sparsity inherent to scATAC-seq data and to increase the number of detectable peak–peak links without substantially altering qualitative results.

The output of this step is a dataframe of peak pairs and their co-accessibility scores, ranging from −1 (anti-correlated accessibility) to 1 (simultaneous accessibility). Since this study focuses on enhancer-mediated regulation, only links with positive co-accessibility scores are retained.

While co-accessibility captures general chromatin coupling, it does not distinguish regulatory interactions involving enhancers from other types of genomic interactions. Consequently, additional biologically informed filtering is required to focus the analysis on enhancer–gene relationships.

#### 3.2.3 Translation and intersection of enhancer–gene associations

To enable direct comparison with GeneHancer, co-accessibility links are translated into enhancer–gene associations. First, peak pairs are filtered to retain only those in which one peak overlaps a *GH*_*el*_ and the other overlaps a gene promoter. Gene coordinates are obtained from the NCBI RefSeq Genes database [O’Leary et al., 2016], and promoter regions are defined as 2,000 bp upstream and 200 bp downstream of the transcription start site, a window commonly adopted in regulatory genomics to capture core promoter accessibility while limiting spurious long-range associations.

Genomic overlaps are computed using pyRanges [Stovner and Sætrom, 2020]. This procedure yields a set of candidate cis-regulatory pairs (*GH*_*el*_, *g*). When multiple peaks map to the same *GH*_*el*_, their co-accessibility scores are averaged to produce a single association score.

The translated co-accessibility-derived associations are then intersected with GeneHancer enhancer– gene links, yielding a final set of conserved associations (**CA**), defined as

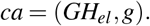

These **CA** represent regulatory interactions that are both supported by curated GeneHancer evidence and reinforced by dataset-specific chromatin accessibility patterns.

### 3.3 Validation of Conserved Associations

A strong argument can be made that both curated regulatory information from GeneHancer and data-driven co-accessibility inference provide meaningful and complementary insights into cis-regulatory interactions when considered independently. GeneHancer associations are supported by extensive cross-source biological evidence, while co-accessibility captures dataset-specific chromatin coupling patterns. However, it remains an open question whether the intersection of these two sources, represented by the conserved associations (**CA**), yields regulatory links that are more biologically relevant than those obtained from either approach alone.

To address this question, this study performs two complementary validation analyses designed to assess the functional relevance of the **CA**. The first analysis focuses on the cell-type specificity of GeneHancer regulatory elements, while the second evaluates the relationship between chromatin accessibility and gene expression. Together, these analyses aim to determine whether **CA** better capture context-specific regulatory mechanisms active in the dataset.

#### 3.3.1 Specificity analysis

The first validation analysis examines the mean cell-type specificity of GeneHancer elements associated with known marker genes. This analysis builds upon the specificity framework introduced in the GRAIGH study [Martini et al., 2023b], which quantifies how selectively a regulatory element is accessible across annotated cell types. Marker genes are expected to exhibit strong and consistent cell-type specificity, and therefore their associated regulatory elements should display similarly constrained accessibility patterns.

For each marker gene and its corresponding annotated cell type, the specificity of the linked *GH*_*el*_ is computed under three distinct scenarios: (1) considering all *GH*_*el*_ associated with the gene in the Gene-Hancer database, (2) restricting the analysis to only *Elite*-status *GH*_*el*_, and (3) considering exclusively the *GH*_*el*_ derived from the conserved associations (**CA**). In each case, specificity values are averaged across all *GH*_*el*_ linked to a given gene, yielding a gene-level specificity score.

By comparing these three scenarios, this analysis evaluates whether the **CA** preferentially retain regulatory elements that are more specific to the biological context of the dataset, relative to the broader and more heterogeneous set of GeneHancer annotations. An increase in specificity for **CA**-derived elements would indicate improved relevance for cell-type–resolved regulatory analysis.

#### 3.3.2 Accessibility–expression correlation

The second validation analysis investigates the functional relationship between chromatin accessibility at *GH*_*el*_ and the expression levels of their associated genes. This analysis integrates matched scATAC-seq and scRNA-seq data from the Multiome dataset to systematically assess whether enhancer accessibility co-occurs with gene expression across cells.

Gene expression values are log-normalized following the standard Scanpy pipeline [Wolf et al., 2018] to ensure comparability across cells. Accessibility profiles for *GH*_*el*_ are extracted from the matrix **GH** _|**GHel** |*×*| *C*|_ generated using GRAIGH (Section 3.2.1). To mitigate the impact of data sparsity and technical noise, the analysis retains only genes with sufficient detectable expression and *GH*_*el*_ that are accessible in at least a predefined fraction of cells.

For each *GH*_*el*_–gene pair, the Jaccard index is computed on binarized accessibility and expression profiles:

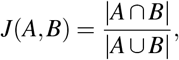

where *A* denotes the set of cells in which a given *GH*_*el*_ is accessible and *B* denotes the set of cells in which the associated gene is expressed. The use of the Jaccard index is motivated by the high sparsity characteristic of scATAC-seq data, which limits the robustness of traditional continuous correlation measures. Although binarization reduces quantitative information, it provides a stable and interpretable metric for assessing co-occurrence between regulatory element activity and gene expression.

The resulting Jaccard values serve as a proxy for functional association between enhancer accessibility and transcriptional output. While this measure does not fully capture the complexity of enhancer–gene regulation, it enables systematic comparison across large sets of associations. In addition to the full set of **CA**, an auxiliary subset is analyzed by restricting to associations supported by co-accessibility scores in the top 75th percentile. This additional filtering step assesses whether stronger chromatin coupling further enhances the correspondence between accessibility and expression, providing insight into the contribution of co-accessibility strength to regulatory relevance.

Together, these validation analyses offer a comprehensive assessment of the biological significance of conserved associations, supporting their use as a refined and context-aware representation of enhancer– gene regulatory interactions.

## 4 RESULTS

### 4.1 Co-accessibility connections and GeneHancer associations

The co-accessibility analysis yielded a total of 1,640,140 peak–peak connections, of which 1,289,311 (80.4%) corresponded to positive co-accessibility scores. In parallel, the GeneHancer database reported 2,408,198 enhancer–gene associations. The size and heterogeneity of both data layers highlighted the fundamental challenge of distinguishing biologically meaningful regulatory interactions from a large background of potentially non-functional connections.

Figure 2 illustrates the progressive filtering of co-accessibility links, GeneHancer associations, and GeneHancer elements (*GH*_*el*_). Among the inferred co-accessibility links, 1,179,560 (73.5%) could be translated into putative *GH*_*el*_–gene associations. This represents a non-trivial result, indicating that a large fraction of co-accessible peak pairs naturally align with candidate cis-regulatory interactions. Of these translated links, 64.9% were retained after intersection with GeneHancer, forming the set of **CA**s. However, when considering the number of unique **CA**s, this figure decreased substantially to 58,912. This reduction is primarily attributable to the fact that individual *GH*_*el*_ frequently overlap multiple peaks. Indeed, while peaks typically spanned 100–400 bp, GeneHancer enhancers often extended over several kilobases [Fishilevich et al., 2017], leading to multiple peak-level links collapsing into a single enhancer–gene association.

**Figure 2.**
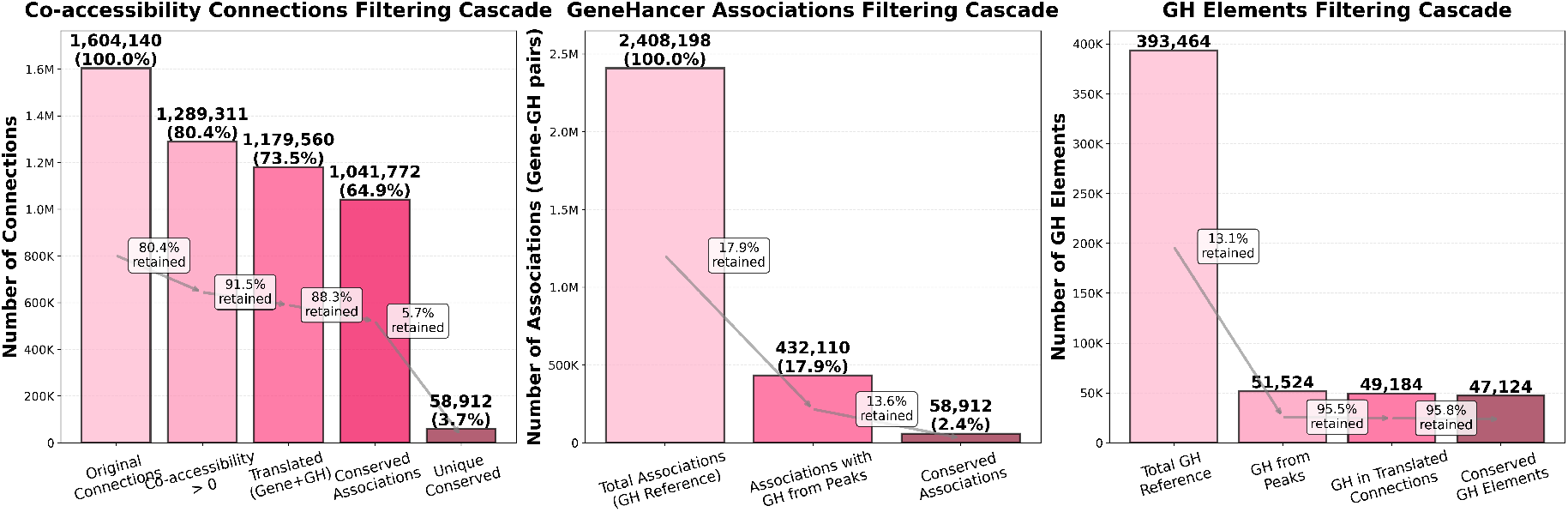
Filtering cascade of co-accessbility connections (left), GeneHancer associations (center), and *GH*_*el*_ (Right).

From the GeneHancer perspective, only 13.1% of all *GH*_*el*_ were accessible in this dataset, and these accessible elements accounted for just 17.9% of all GeneHancer-reported enhancer–gene associations (Figure 2). This observation underscores that the majority of GeneHancer annotations correspond to regulatory elements that were inactive in PBMCs and would therefore have introduced substantial noise and non-specificity if used without contextual filtering. The intersection of GeneHancer with co-accessibility further reduced the set to only 2.4% of all Gene-Hancer associations. Although numerically small, this subset was enriched for interactions that were both supported by prior biological evidence and active in the specific cellular context under investigation.

Moreover, the **CA**s represent only 13.6% of the associations involving peak-overlapping *GH*_*el*_. This reflects the fact that many *GH*_*el*_ were linked to genes not expressed in the dataset, or were associated with multiple genes while regulating only a subset in a given biological context. Notably, the number of *GH*_*el*_ retained after filtering was similar whether one considered all accessible *GH*_*el*_ or only those participating in a **CA**. This suggests that most regulatory elements relevant to the dataset’s transcriptional landscape were also captured within the conserved set, providing further support that integrating Gene-Hancer with co-accessibility effectively prioritized biologically active enhancers.

At the same time, co-accessibility alone did not constitute a definitive indicator of functional regulation. Owing to its purely data-driven nature, co-accessibility reflected coordinated chromatin accessibility rather than direct regulatory causality. Consequently, many inferred links may have represented indirect associations, structural chromatin effects, or technical artifacts, rather than enhancer–gene interactions. Co-accessibility therefore benefited substantially from integration with curated regulatory knowledge to improve biological interpretability.

Overall, these results highlight the complementarity between curated, cross-dataset regulatory annotations and data-driven epigenomic evidence. Gene-Hancer provides broad functional context, while co-accessibility captures the sample-specific regulatory architecture. Their intersection isolated a focused, high-confidence set of enhancer–gene associations that is both biologically grounded and context-aware, offering a robust foundation for downstream regulatory analyses and interpretation of cell identity programs.

### 4.2 The conserved associations shows higher specificity and correlation

After identifying the conserved associations, we assessed whether this refined set retains and enhances biologically relevant regulatory signals. The analysis focused on well-established marker genes of the major PBMC cell types (i.e., CD4^+^ and CD8^+^ T cells, monocytes, B cells, and natural killer (NK) cells) given the expectation that these genes are governed by highly cell-type–specific regulatory programs.

Figure 3 shows the heatmap of specificity values across three sets of GeneHancer elements: all GeneHancer-associated elements, *Elite* GeneHancer elements, and elements derived from the conserved associations. Across all marker genes and cell types, *GH*_*el*_ from **CA**s consistently exhibited markedly higher specificity than those drawn directly from the full GeneHancer database. This result indicates that relying solely on unfiltered GeneHancer annotations introduces non-specific regulatory information that dilutes genuine cell-type relevance.

**Figure 3.**
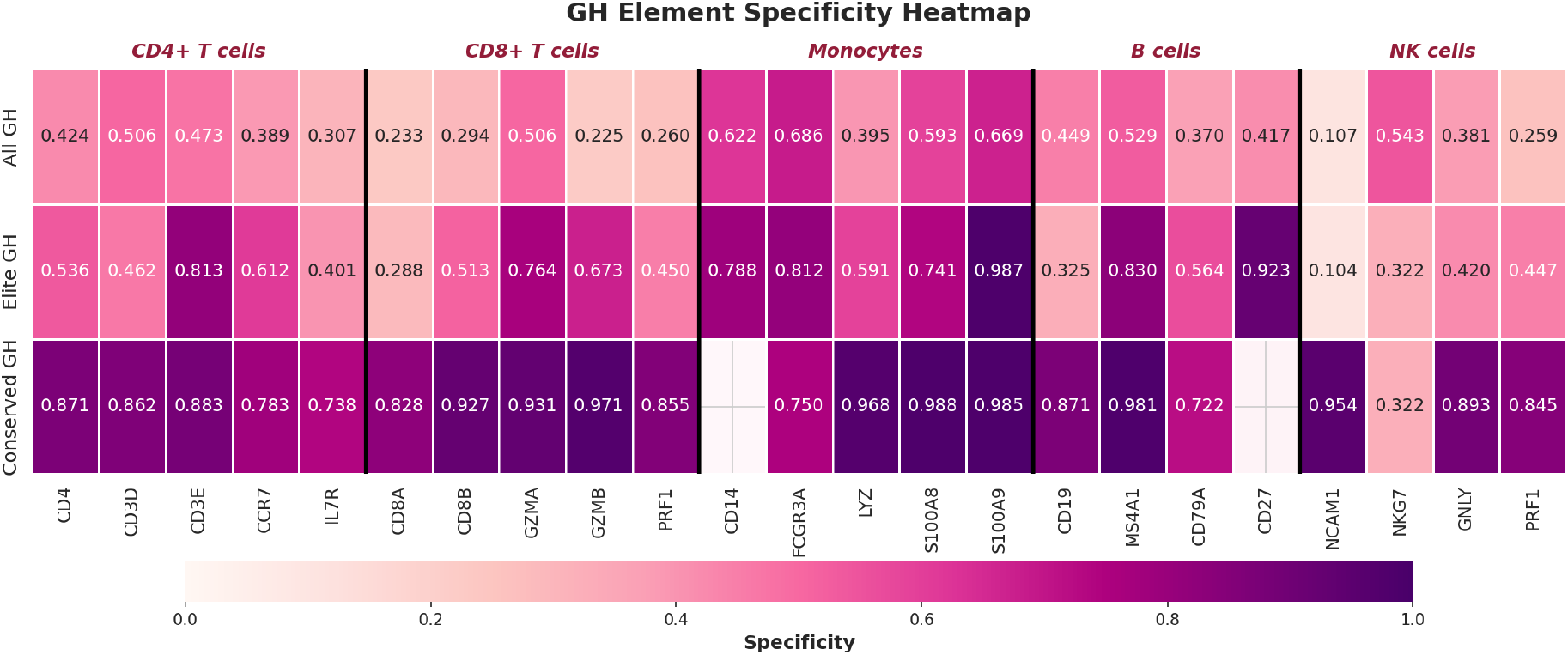
Heatmap of the mean specificity of *GH*_*el*_ associated with known marker genes under different association selection strategies.

Restricting the analysis to GeneHancer *Elite* associations, i.e., those supported by multiple independent evidence sources, improved specificity relative to the full database but still did not outperform the conserved set. While elite associations partially mitigate noise, they remained less specific than the dataset-informed **CA**, underscoring the importance of incorporating epigenomic evidence rather than relying exclusively on cross-dataset curation.

For two genes, *CD14* and *CD27*, no conserved associations were identified. In the case of *CD14*, this absence is likely attributable to the presence of regulatory elements shared among multiple nearby genes, which can obscure gene-specific regulatory contributions. This observation highlights an intrinsic limitation of co-accessibility-based approaches: shared enhancers may distribute their signal across several potential targets, complicating gene-level assignment. Future extensions incorporating cell-type–specific or single-cell–resolved co-accessibility estimation may help alleviate such confounding effects.

Accessibility–expression correlation analyses provided an additional layer of validation (Figure 4). Although these correlations were generally weaker and sparser due to the distinct characteristics of the two modalities, **CA**s consistently exhibited equal or higher correlation values compared to the other association sets. Notably, *IL7R* and *MS4A1* displayed substantially stronger and more coherent correlation patterns within the conserved set, accompanied by higher Jaccard indices. For *MS4A1*, the mean co-accessibility score of the associated *GH*_*el*_ was 0.5141, exceeding the 75th percentile of all inferred connections (0.2233), indicating a robust and biologically plausible regulatory relationship. Such strong associations not only support downstream dataset-specific analyses but may also serve as candidates for cross-validation and refinement of existing regulatory databases.

**Figure 4.**
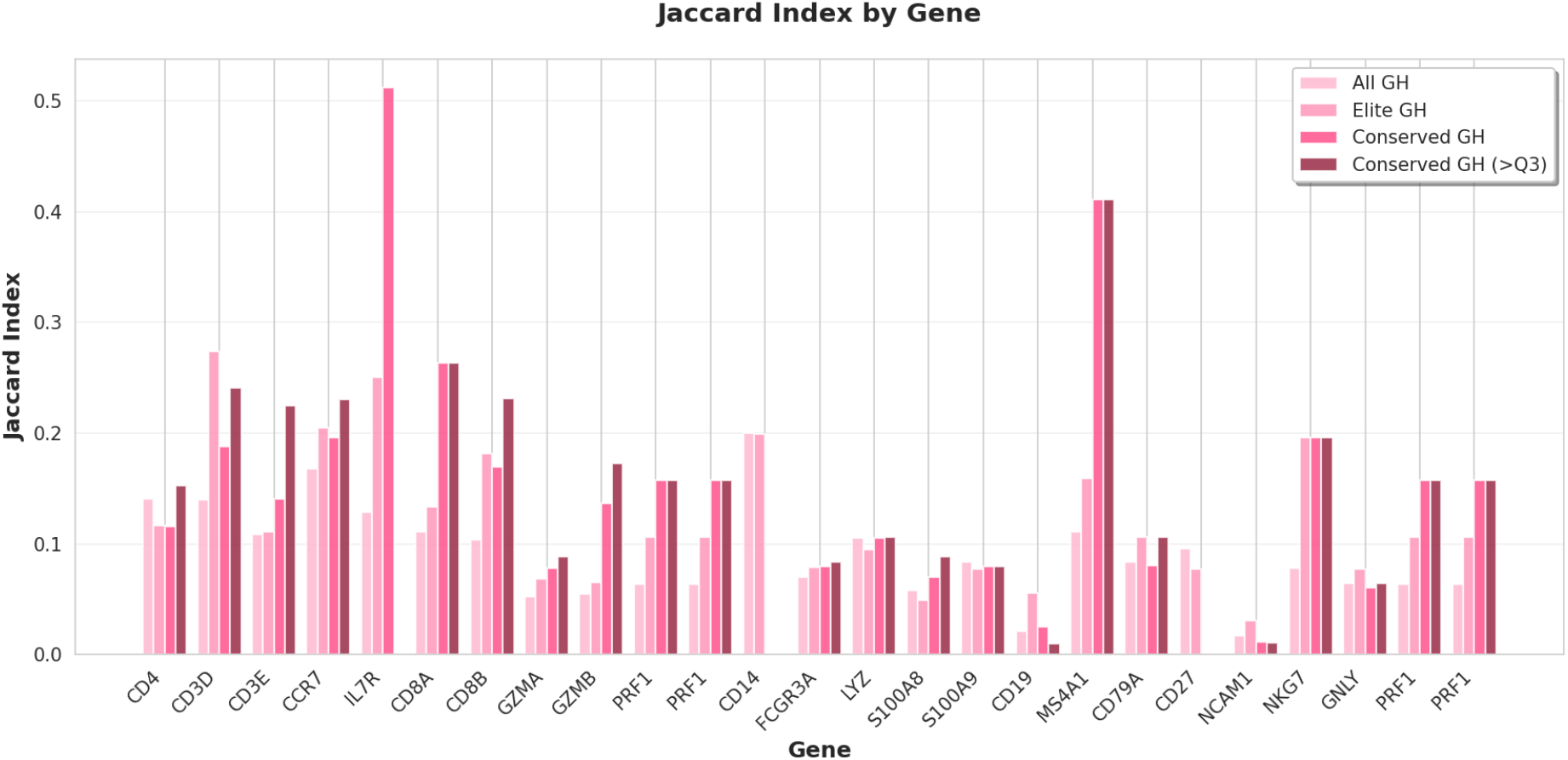
Barplot of the mean Jaccard Index of *GH*_*el*_ associated with known marker genes. The plot shows also the **CA** filtered over the 75 percentile.

Consistent with this observation, analysis of the subset of **CA** supported by co-accessibility scores above the 75th percentile revealed that additional filtering either preserved correlation metrics, indicating that highly co-accessible links were already captured by the initial intersection, or further increased them. This trend suggests that stronger chromatin coupling is generally associated with improved concordance between accessibility and expression.

Several limitations should be acknowledged. First, the analysis is demonstrated on a human PBMC dataset, and regulatory interactions identified here may not generalize to other tissues or developmental contexts without reapplication of the framework to additional datasets. Second, the approach depends on the quality and coverage of scATAC-seq data; sparse accessibility profiles may lead to false negatives, particularly for weakly active or shared enhancers. Third, co-accessibility reflects coordinated chromatin accessibility rather than direct regulatory causality, and some functional interactions may remain undetected.

Finally, although the use of metacells improves robustness, co-accessibility inference remains computationally demanding for very large datasets. Future work will focus on extending the framework to additional biological systems, incorporating cell-type–specific co-accessibility estimation, and integrating complementary evidence such as chromatin conformation or perturbation data to further improve enhancer–gene assignment.

Taken together, these results demonstrate that **CA**s represent a refined and biologically coherent subset of enhancer–gene associations. They capture enhanced regulatory specificity, reflect dataset-driven epigenomic structure, and show improved agreement with gene expression patterns, supporting their utility for downstream regulatory analyses and the interpretation of cell identity programs.

## 5 CONCLUSIONS

Integrating curated enhancer–gene annotations with data-driven co-accessibility patterns provides an effective strategy for refining cis-regulatory interactions in a dataset-specific manner. While GeneHancer offers a broad, functionally supported compendium of regulatory relationships, our results showed that only a small fraction of these associations is active in the PBMC dataset analyzed. Intersecting GeneHancer annotations with co-accessibility networks derived from scATAC-seq data yielded a focused set of enhancer–gene associations exhibiting higher cell-type specificity, stronger concordance between chromatin accessibility and gene expression, and clearer biological relevance than those obtained from either the full or the *Elite* GeneHancer sets alone.

These findings highlight the complementary strengths of curated regulatory databases and single-cell epigenomic evidence. Curated resources contribute robust cross-study validation, whereas co-accessibility captures the regulatory architecture active within a specific biological context. Their integration reduces redundancy, filters out inactive regulatory elements, and prioritizes enhancer–gene interactions supported by both prior knowledge and dataset-specific chromatin signals. The refined associations produced by this approach may also inform future improvements in enhancer annotation by identifying regulatory relationships that are consistently supported across orthogonal evidence sources. More broadly, this framework underscores the value of combining model-driven resources with data-driven analyses to more accurately delineate the regulatory landscape of the non-coding genome and to support downstream investigations of gene regulation and cell identity.

## 6 Code Availability

GeneCard allows direct download of the older database 2017 version https://www.genecards.org/GeneHancer Version 4-4, but it is possible to request the access to the latest versions from the online platform https://www.genecards.org/Guide/DatasetRequest. The 10X genomics dataset is freely available at https://www.10xgenomics.com/resources/datasets/10k-human-pbmcs-atac-v2-chromium-controller-2-standard. All the code employed in this study is publicly available on the GitHub repository at https://github.com/smilies-polito/GH-Co-Accessibility.git, with a dedicated singularity container and an automatic SnakeMake [Mölder et al., 2021] pipeline, to ensure full reproducibility.

## ACKNOWLEDGEMENTS

The authors acknowledge the use of GPT-4.0 and GPT-4.0 with optimization (GPT-4o and GPT-4o1) for language and text refinement during manuscript preparation. Authors employed these tools to enhance clarity, coherence, and readability, ensuring high-quality presentation of the research findings. The scientific content, data analysis, and interpretations remain the authors’ sole responsibility. This work was supported by project SERICS (PE00000014) under the MUR National Recovery and Resilience Plan funded by the European Union.

